# Simplifying the combined use of CRISPR-Cas9 and Cre-loxP technologies for the efficient generation of targeted conditional gene knockouts in mammalian cells

**DOI:** 10.1101/236679

**Authors:** Tzahi Noiman, Chaim Kahana

**Affiliations:** Department of Molecular Genetics, the Weizmann Institute of Science, Rehovot 76199, Israel

## Abstract

Gene knockout technologies have contributed fundamentally to our understanding of the cellular functions of various genes. Two prevalent systems used for the efficient elimination of the expression of specific genes are the Cre-LoxP system and the CRISPR-Cas9 system. Here we present a simple method that combines the use of CRISPR-Cas9 and Cre-loxP for the conditional deletion of essential genes in mammalian cells. First, an inducible Cre recombinase is stably expressed in the cells. Next CRISPR-Cas9 is used to knockout an essential gene, whose function is complemented by stable expression of a FLAG-tagged version of the same protein encoded from a floxed transcription unit containing silent mutations, making it refractory to the CRISPR-Cas9 guide. This FLAG-tagged protein can be deleted by activating the expressed Cre protein enabling evaluation of the cellular consequences of its deletion. We have further used this system to evaluate potential mutants of the tested gene.

## INTRODUCTION

Gene knockdown/knockout technology has contributed fundamentally to our understanding of the cellular functions of various genes. Knockdown is achieved by using siRNA that can be used transiently or shRNA that can also be stably expressed in target cells (1–3). However, since these methods reduce, but do not eliminate the expression of target genes, the biological consequences on the affected cells are not always complete. Efficient elimination of gene expression is therefore required to obtain a comprehensive assessment of the cellular role of a given gene. Two main systems have manifested themselves as the leading technologies for achieving this goal. The first is the Cre-LoxP technology (4, 5), and the second is the CRISPR-Cas9 system, which revolutionized genome editing capability (6).

Cre is a 38-kDa site specific DNA recombinase that recognizes a 34-bp sequence denoted loxP, resulting in intra and inter molecular recombination between two loxP sites depending on their orientation and location (7). The loxP sites, composed of an 8-bp non-palindromic core flanked by two 13-bp inverted repeats, are usually placed in the desired locations and orientations in the desired DNA sequence via homologous recombination using a construct containing selectable markers (8). When two LoxP sites flank a sequence and are directly repeated, Cre excises the entire DNA segment located between them.

CRISPR-Cas9 is a novel and currently the most efficient genome editing tool. The widely used CRISPR-Cas9 system is comprised of two components; a guiding RNA, referred to as gRNA, and an endonuclease and helicase enzyme called Cas9, capable of cutting double stranded DNA at a specific genomic location, depending on the targeting sequence presented in the gRNA sequence. The single guide RNA is composed of 20 nucleotides, homologous to the genomic site to be edited, and an RNA scaffold portion that facilitates Cas9 binding to the DNA site to generate the double-strand breaks (DSBs) at the target genomic site (9–11). The DBSs are usually repaired by the Non-Homologous End-Joining (NHEJ) pathway that frequently generates deletions or insertions, seldom resulting in abolishing the expression of the tested protein. Variants of Cas9 are still being developed to enhance specificity and minimize off-target editing (12–14).

A less efficient, high fidelity repair pathway employed by cells is the Homology Directed Repair (HDR) pathway. This high fidelity repair pathway can be exploited by mutating Cas9 at one of its catalytic domains to create a nickase variant, termed Cas9 nickase, capable of generating single-stranded DNA breaks that together with a DNA “repair template”, allow repair to proceed by HDR rather than NHEJ, leading to a more precise gene editing or to the addition of sequences to endogenous DNA, depending on the template used (15). The process is limited by relatively low efficiency, and several methods have been developed in order to increase efficiency (16) but not without possible negative effects(17).

While both methods are suitable for the investigation of many genes, inducible systems are required for the investigation of essential genes and for investigating the temporal importance of genes during development. To achieve inducibility, Cre is fused to a mutated hormone-binding domain of the estrogen receptor, generating a chimeric Cre recombinase, termed CreER, that is activated by synthetic estrogen receptor ligands (18–21). CRISPR-Cas9 inducible systems were also developed, which rely either on an inducible promoter or on the activation of a CRISPR-Cas9 estrogen receptor chimera (22–25). However, expression of Cas9 from inducible promoters can be leaky, (26)(27), and the introduction of LoxP sites into a specific genomic location requires extensive constructions and manipulations, seldom with varying efficiencies (28–30).

Here, we present a simple method that combines the CRISPR-Cas9 and the Cre-loxP systems. This method is particularly useful for the conditional deletion of essential genes in mammalian tissue cultured cells and can further be exploited to identify mutants of tested genes.

## MATERIAL AND METHODS

### Plasmid constructs

ER^T2^CreER^T2^ Was PCR amplified using pCAG-ERT2CreERT2 (Addgene # 13777) as a template and was cloned between Xhol and Notl sites of the bicistronic pEFIRES vector (31) harboring a selectable marker conferring resistance to G418.

To generate pX330-5A1, a 19-nt guide sequence (Table 1) targeting exon 1 of the mouse eIF5A1 gene was ligated into a pX330 hSpCas9 plasmid (Addgene #42230).

**Table 1.**
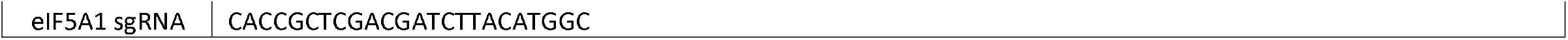
sgRNAs sequences used in the paper.

pE-eIF5A1smut, containing a FLAG-tagged eIF5A1 that is refractory to the guide RNA, was constructed in two steps; first, eIF5A1 cDNA was amplified by PCR using a 5’ primer containing a LoxP site and appending a FLAG tag to the N-terminus of eIF5A1 and a 3’ primer containing a LoxP site. The resulting fragment was cloned into the bicistronic pEFIRES vector containing a selectable marker conferring puromycin resistance (Table 2). Next, three silent mutations were introduced at the site of recognition by the guide RNA to make it refractory to editing by the CRISPR-Cas9 (Table 2).

**Table 2.**
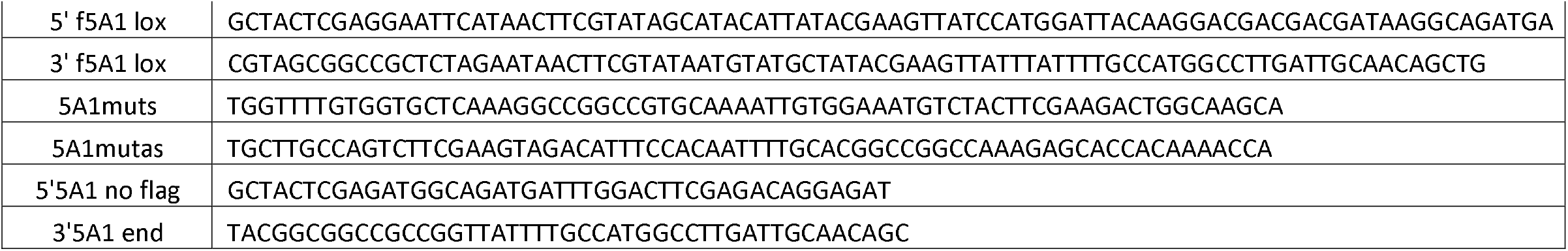
Primers used in the paper.

Wild type eIF5A1(pWT-5A1) and a K50R mutant (pK50R-5A1) that cannot undergo hypusination and harbouring mutations that makes them refractory to recognition by the gRNA were PCR amplified using eIF5A1 specific primers and cloned into a bicistronic pEFIRES vector harbouring a selectable marker conferring hygromycin resistance.

### Cell culture

NIH3T3 mouse fibroblasts were grown in Dulbecco’s modified Eagle’s medium (Invitrogen) supplemented with 10% (v/v) fetal bovine serum, 100 units/ml penicillin, and 100 μg/ml streptomycin (Biological Industries).

NIH3T3 cells were transfected using Lipofectamine 2000 (Invitrogen) following the manufacturer’s instructions. Twenty four hours post transfection the cells were split and subjected to the indicated selections. When validation of a successful knockout was required, individual foci were selected, expanded and subjected to western blot analysis using an anti eIF5A1 antibody. Positively identified clones were subjected to single cell cloning.

### Immunoblot Analysis

Cellular extracts were prepared by lysing cells in RIPA buffer (50 mm Tris-HCl pH 8, 150 mm KCl, 1.0% Nonidet P-40 (IGEPAL), 0.5% sodium deoxycholate, 0.1% SDS) supplemented with protease inhibitors cocktails (Sigma). Equal portions of protein were resolved by electrophoresis in SDS-polyacrylamide gel, blotted to a nitrocellulose membrane and incubated with the indicated antibodies followed by horseradish peroxidase-conjugated anti-IgG antibodies. The antibodies used were: mouse mAb anti-eIF5A1 (BD Transduction Laboratories, #611976), mAb anti-FLAG M2 (Sigma, # F1804). Signals were developed using “EZ-ECL” (Biological Industries), and the membranes were analysed using ImageQuant LAS4000 luminescent image analyser (General Electric).

## RESULTS

### Establishment of a cell line expressing inducible Cre

The first step in the establishment of the inducible gene knockout system was the cloning of a variant of CreER^T2^, ER^T2^CreER^T2^ [currently the most tightly regulated version of CreER (32, 33)], which allows for even tighter regulation compared to CreER^T2^ (34). ER^T2^CreER^T2^ was cloned as the first open reading frame (ORF) of the bicistronic vector pEFIRES (35), and the resulting construct was transfected into NIH3T3 mouse fibroblasts. Since the second ORF of the bicistronic transcript encodes resistance to G418, the transfected cells were selected for growth in the presence of G418 (500ug/ml). G418 resistant cells were selected and tested for functionality and inducibility of the ER^T2^CreER^T2^ by transient transfection with the Stoplight construct (36), and induction using 4-Hydroxytamoxifen(4-OHT).

### CRISPR-Cas9 mediated knockout

Following the establishment of cells stably expressing ER^T2^CreER^T2^, we next set up to knockout the gene encoding eIF5A1 (37), using CRISPR-Cas9 (38). Deskgen’s Guide Picker (39) was used to identify optimal target site. Oligonucleotides representing the single-guide RNA (sgRNA) were cloned in the pX330-U6-Chimeric_BB-CBh-hSpCas9 plasmid (40). Since eIF5A1 is an essential gene (41), the resulting Cas9 construct was co-transfected into the ER^T2^CreER^T2^ expressing NIH3T3 cells together with a bicistronic construct expressing a floxed, flag tagged eIF5A1 containing a silent mutation making it refractory to the guide RNA. The puromycin resistance encoded as a second ORF of the bicistronic transcript enabled the isolation of individual foci. Western blot analysis performed on cellular extracts prepared from cells representing individual foci identified clones that lost their endogenous eIF5A1 but express the FLAG-tagged eIF5A1 expressed from the floxed allele (Figure 1A).

**Figure 1.**
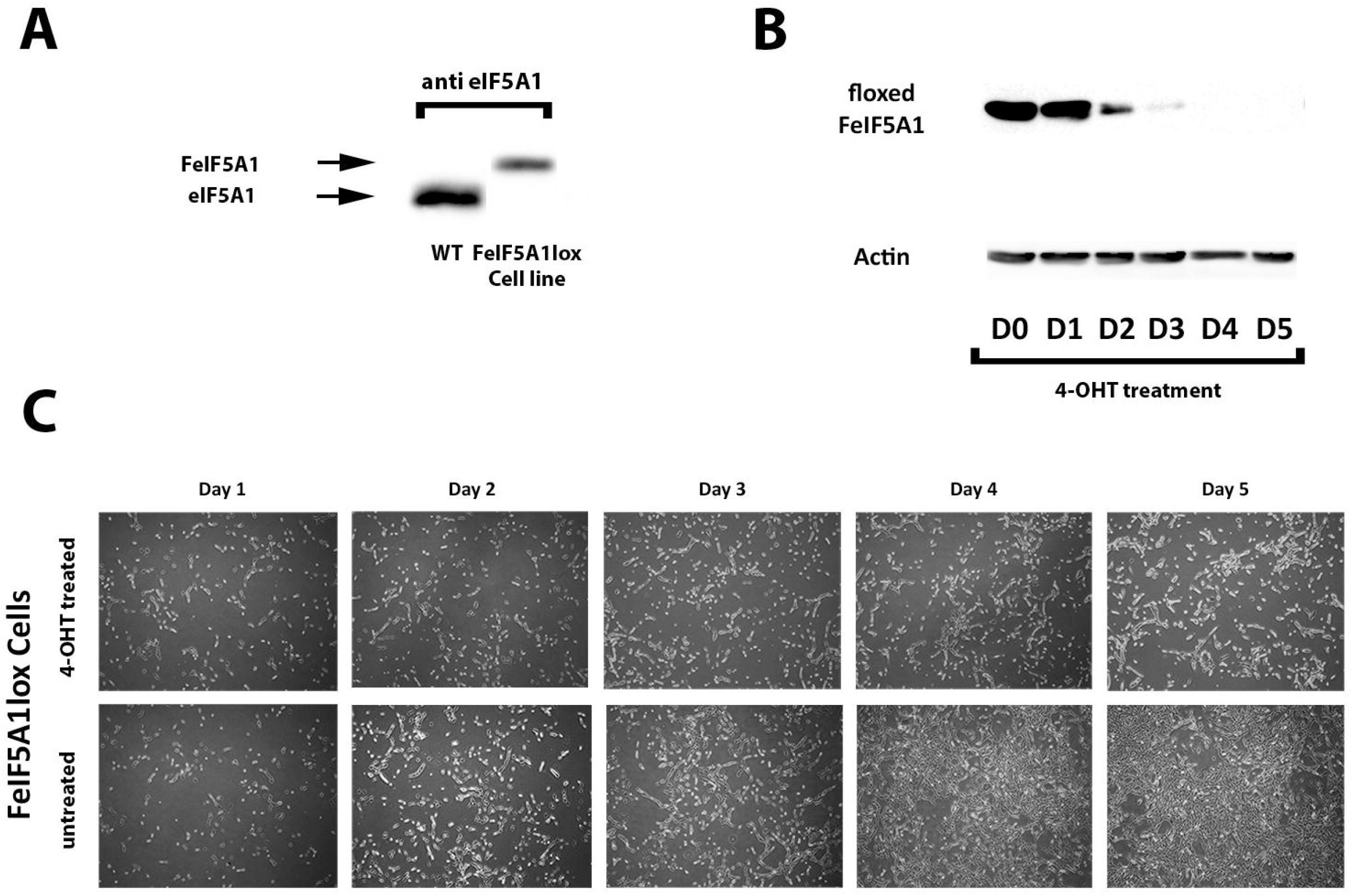
Establishment of an inducible knockout cell line using a combined Cre/Lox and CRISPR/Cas9 systems. (A) Wild-type NIH3T3 cells stably expressing ER^T2^CreER^T2^ Were co-transfected with a pX330 Cas9 construct containing a sgRNA against exon 1 of the eIF5A1 gene together with a bicistronic vector expressing from its first ORF a FLAG-tagged floxed version of eIF5A1 harboring a silent mutations making it refractory to Cas9 editing of eIF5A1, and puromycin resistance from its second ORF. FeIF5A1lox represents the cell line that does not express endogenous eIF5A1 while ectopically expressing a FLAG-tagged eIF5A1 protein encoded from a floxed allele. The position of the endogenous WT eIF5A1 and of the ectopically expressed FLAG-tagged mutant proteins are indicated by arrows. (B) FeIF5A1lox cells were treated with 4-OHT to induce Cre-Lox recombination. Cellular extracts were prepared at the indicated times and subjected to western blot analysis using an anti-FLAG antibody. An anti Actin antibody was used as control. (C) FeIF5A1lox cells were treated with 4-OHT to induce Cre-Lox recombination. Cells were documented at the indicated time points following 4-OHT induction.

### Induction of Cre-mediated deletion of the ectopically expressed FLAG-tagged eIF5A1

Since the FLAG-tagged eIF5A1 encoded from the bicistronic plasmid is also flanked with unidirectional loxP sequences, it can be deleted by activating the ER^T2^CreER^T2^ encoded in these cells. To induce deletion of the FLAG-tagged floxed eIF5A1 transgene, the cells were treated with 1μM 4-OHT. As can be seen in figure 1B, the levels of the FLAG-tagged eIF5A1 declined gradually, becoming very low already at day 2, almost undetected at day 3 and completely absent from day 4 and on. The depletion of eIF5A1 was accompanied by cessation of cellular proliferation noted between day 2 and 4 (Figure 1C).

### Screening for mutant proteins

We next set out to test the possibility that the inducible knockout system we present here can also be used for screening for potential mutants of the tested proteins. For this purpose we cloned a wild-type (pWT-5A1) and a non-hypusinated mutant of eIF5A1 (pK50R-5A1) into a bicistronic vector encoding at its second ORF hygromycin resistance as a selectable marker. Hypusination is a unique post translational modification of eIF5A proteins demonstrated to be essential for their functionality (41–44). An empty vector was used as negative control (Figure 2B). Following transfection, cells growing in the presence of hygromycin were selected and induced by the addition of 4-OHT to the growth medium. As expected, decline in the level of the FLAG-tagged eIF5A1 encoded by the floxed allele was observed, whereas the expressed wild type and non-hypusinated mutant remained unaffected (Figure 2A). Concomitantly, cells that were stably transfected with the wild-type encoding construct remained proliferating, while those transfected with an empty vector or with the vector encoding the non-hypusinated mutant became growth arrested (Figure 2B).

**Figure 2.**
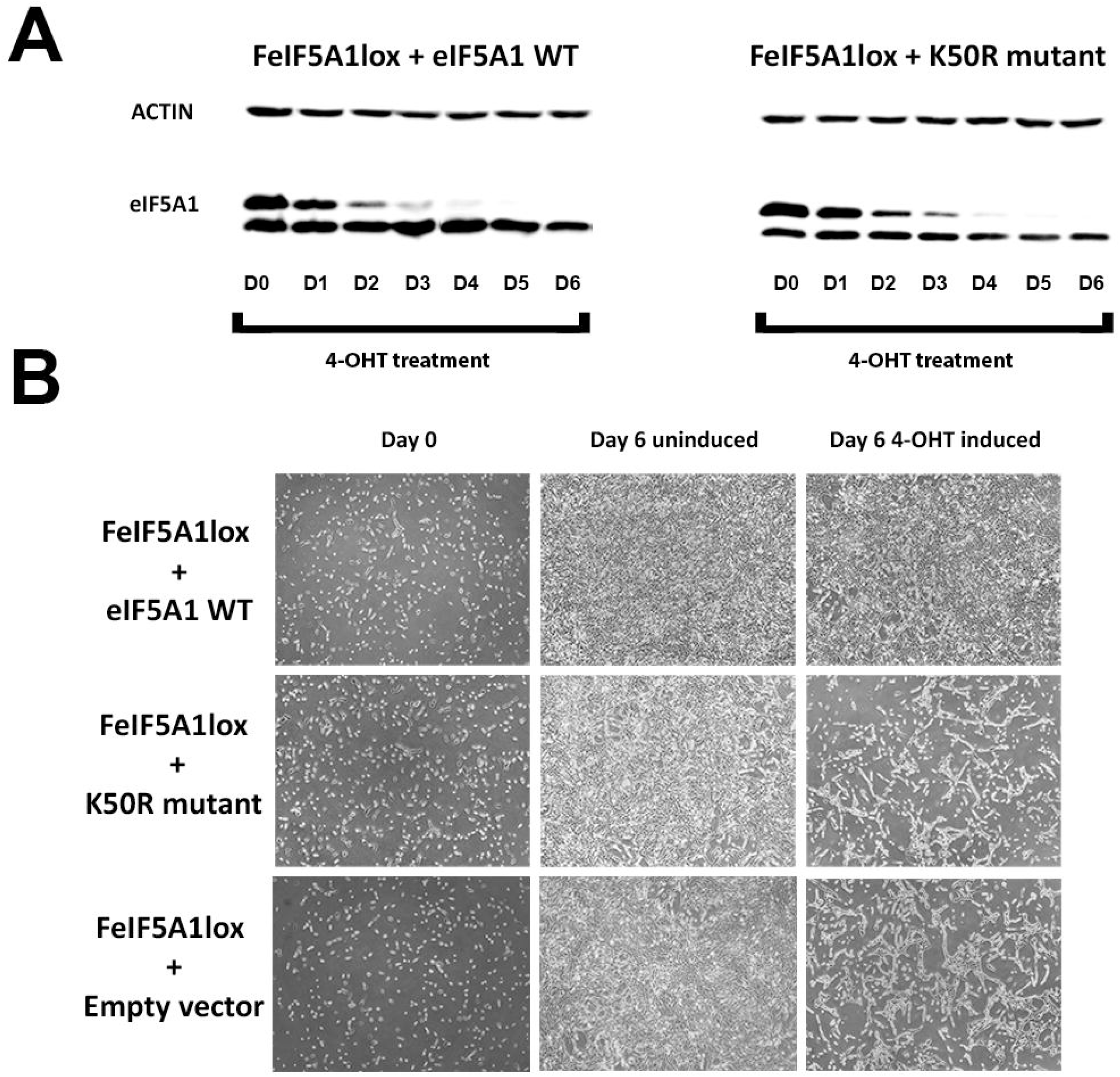
Repurposing FeIF5A1lox cells for screening of mutant eIF5A1 proteins. FeIF5A1lox cells were stably transfected with a bicistronic construct encoding either wild type eIF5A1 (eIF5A1 WT), or a non-hypusinated eIF5A1 mutant (K50R mutant), from its first ORF, and a hygromycin B resistance from its second ORF. Representative clones were induced using 4-OHT to induce Cre-Lox recombination of the floxed, FLAG-tagged eIF5A1. (A) Western blot analysis of cellular extracts prepared at the indicated times following 4-OHT induction. (B), cells harboring an empty vector or constructs encoding the WT or mutant proteins were treated with 4-OHT to induce Cre-Lox recombination. Cells were documented at the indicated time points following 4-OHT induction.

## DISCUSSION

We present here a simple and efficient method that enables the generation of conditional knockouts of essential genes in mammalian cells (Figure 3). This method combines two highly efficient platforms, the CRISPR-Cas9 and the Cre-loxP systems. In this system, conditional knockout is manifested in two simple steps. In the first step, ER^T2^CreER^T2^ is stably expressed in the target cells. This step can be performed as the starting point in any cell line before any downstream steps are taken. In the second step, two vectors are utilized simultaneously: a SpCas9 expressing vector with a gRNA designed to knockout the tested gene, and a bicistronic vector encoding a modified cDNA encoding a FLAG-tagged version of the protein encoded by the gene to be deleted. The cDNA is modified in two ways; it contains a silent mutation which makes it refractory to the CRISPR-Cas9 mediated DNA editing, and it is flanked by two LoxP sites added by PCR. The bicistronic transcript provides two functions. The puromycin resistance encoded by the second ORF enables the selection of foci, while the flag-tagged protein encoded by the first ORF provides a “back-up” that enables the deletion of the two genomic alleles encoding the essential protein. Successful deletion is verified by western blot analysis, relying on the size difference between the endogenous protein and the ectopically expressed FLAG-tagged protein, or by genomic analysis. ER^T2^CreER^T2^ can then be activated by the addition of 4-OHT to the growth medium, resulting in the deletion of the ectopically expressed FLAG-tagged protein, enabling the testing of the cellular role of the deleted protein.

**Figure 3.**
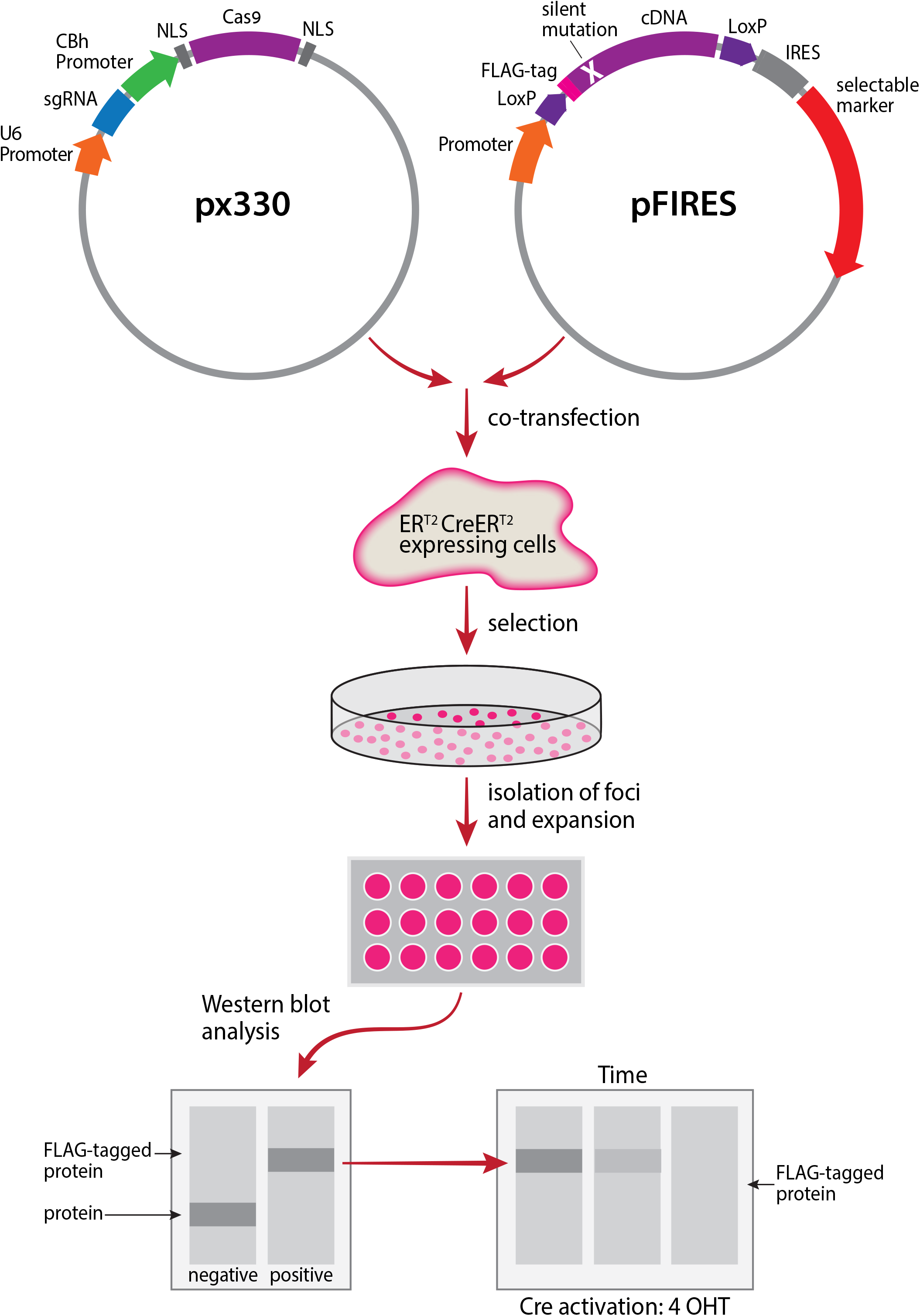
Schematic representation of experimental design. NIH3T3 cells are transfected with a bicistronic construct encoding the inducible ER^T2^CreER^T2^ protein. Stable transfectants are selected for growth in the presence of G418. The resulting cells are then cotransfected with a plasmid encoding a sgRNA targeting the gene of interest and Cas9 together with a construct expressing a FLAG-tagged version of the gene to be deleted that contains silent mutations making it refractory to recognition by the gRNA. The DNA fragment encoding this protein is flanked by two LoxP sites. Individual foci are isolated, expanded and tested by western blot analysis. Foci lacking the wild-type protein, while expressing its FLAG-tagged version are then selected (by their ability to grow in the presence of puromycin), for further studies. The encoded floxed FLAG-tagged protein can be depleted from the cells by activating the ER^T2^CreER^T2^, enabling evaluation of the cellular role of the tested protein. Another layer of analysis can be performed by expression of mutants of the analyzed protein via transfected bicistronic vector providing hygromycin resistance. The functionality of these mutants is evaluated after depleting the FLAG-tagged wild type protein by activating the ER^T2^CreER^T2^.

Our method has several advantages compared to other methods of constructing inducible knockouts (45–47). It is very simple and easy to be implemented since it is based on simple vectors without requiring complex genomic manipulation. The LoxP sites are introduced by simple PCR reaction into a plasmid without requiring their complicated introduction into the relevant genomic locus via homologous recombination. There is also no requirement for intermediate manipulation for the removal of the selectable marker used for the genomic introduction of the LoxP sites. The system is robust, tightly regulated and results in complete depletion of the tested protein. A further advantage is an additional layer of analysis that permits testing and screening potential mutants of the deleted protein, achieved by cloning cDNA encoding for specific mutants into a third bi-cistronic construct that provides resistance to hygromycin. Upon inducing deletion of the floxed supporting wild-type protein, the ability of the mutants to substitute the wild-type protein can be tested. This allows for a streamlined exploration of mutants, which can be further improved using a random barcoded library of mutants (48–50).

## Acknowledgments

Chaim Kahana is the incumbent of the Jules J. Mallon Professorial chair in Biochemistry.

## FUNDING

This work was supported by grant 855/15 from the Israel Academy of Science and Humanities

## CONFLICT OF INTEREST

The Authors declare that there are no competing interests associated with this manuscript.

